# Drug prescription pattern in European neonatal units

**DOI:** 10.1101/463240

**Authors:** Inge Mesek, Georgi Nellis, Jana Lass, Tuuli Metsvaht, Heili Varendi, Helle Visk, Mark A. Turner, Anthony J. Nunn, Jennifer Duncan, Irja Lutsar

**Author notes:** **Corresponding author:** Inge Mesek, Department of Microbiology, University of Tartu.

## Abstract

**Background:** Hospitalized neonates receive the highest number of drugs compared to all other age groups, but consumption rates vary between studies depending on patient characteristics and local practices. There are no large scale international studies on drug use in neonatal units. We aimed to describe drug use in European neonatal units and characterize its associations with geographic region and gestational age (GA).

**Methods:** A one-day point prevalence study (PPS) was performed as part of the European Study of Neonatal Exposure to Excipients (ESNEE) from January to June 2012. All neonatal prescriptions and demographic data were registered in a web-based database. The impact of GA and region on prescription rate were analyzed with logistic regression.

**Results:** In total, 21 European countries with 89 neonatal units participated. Altogether 2173 prescriptions given to 726 neonates were registered. The 10 drugs with the highest prescription rate were multivitamins, vitamin D, caffeine, gentamicin, amino acids for parenteral nutrition, phytomenadione, ampicillin, benzylpenicillin, fat emulsion for parenteral nutrition and probiotics. The six most commonly prescribed ATC groups (alimentary tract and metabolism, blood and blood-forming organs, systemic anti-infectives, nervous, respiratory and cardiovascular system) covered 98% of prescriptions. GA significantly affected the use of all commonly used drug groups. Geographic region influenced the use of alimentary tract and metabolism, blood and blood-forming organs, systemic anti-infectives, nervous and respiratory system drugs.

**Conclusions:** While GA-dependent differences in neonatal drug use were expected, regional variations (except for systemic anti-infectives) indicate a need for cooperation in developing harmonized evidence-based guidelines and suggest priorities for collaborative work.

## BACKGROUND

Medicines play a pivotal role in improving neonatal health and reducing mortality, and thus are widely used in neonatal units. A median of 3 to 11 drugs per neonate are given depending on the setting and gestational age (GA) [1,2]. Most drugs administered to neonates are either off-label or unlicensed and the exposure of off-label/unlicensed drugs is the highest among preterms [1]. The use of drugs depends on underlying conditions, which are often associated with GA, drugs availability in a country and presence of evidence-based guidelines. Traditions and expert opinions could play a role as well [3,4].

Drug use among neonates has been poorly studied. The most comprehensive studies conducted in the United States based on national datasets showed that antibiotics – ampicillin and gentamicin were the most commonly used medicines [5,6]. European studies have covered single countries, single centers or only selected therapeutic groups like antibiotics [7,8]. Due to methodological variabilities between-country comparisons are complicated. Still, significant geographical differences in drug use have been described [9]. Describing drug use patterns is crucial in obtaining a comprehensive picture of the present situation and identifying priority areas for research.

The European Study of Neonatal Exposure to Excipients (ESNEE) was a pan-European project that aimed to describe the use of pharmaceutical excipients in neonatal drugs. For this purpose data on drug use in neonatal units were collected as previously described [10]. In this substudy of ESNEE we aimed to describe the use of drugs in European neonatal units and explore how geographic region and GA influence their consumption. We hypothesized that GA will influence drug use regardless of region because underlying conditions depend on GA, and drug use should not depend on geographic region.

## METHODS

A multicenter, single-day point prevalence study (PPS) was performed as detailed elsewhere [2,7,10]. Briefly, all units with >50% neonatal admissions in 27 European Union countries plus Iceland, Norway, Serbia and Switzerland, were invited to participate. Data collection was performed in a web-based database within one day, chosen by the unit, within three fixed two-week study periods from January 01^st^ to June 30^th^, 2012. The study sites were divided according to the United Nations Statistics department [11] European geographical division (East, North, South, West). For participating units level of care (I, II and III) [12] and hospital teaching status (teaching/non-teaching) were registered.

All neonates aged ≤28 days present at 8am in the neonatal unit and receiving prescriptions on the study day were included and demographic data was recorded. Neonates were categorized based on GA to extremely preterm (22–27 weeks), very preterm (28–31 weeks), late preterm (32–36 weeks) and full-term (≥37 weeks) [13].

All prescriptions excluding blood products, glucose and electrolyte solutions, vaccines, nursery care topical agents, herbal medicines and enteral nutrition including breast milk fortifiers, were collected. For every drug trade name, active pharmaceutical ingredient (API), dose and route of administration were registered.

The prescriptions were analyzed based on the World Health Organization Anatomical Therapeutic Chemical (ATC) classification system [14] according to the level 1 (main anatomical), 3 (pharmacological subgroup) and 5 (chemical substance). Prescription rates (number of prescriptions per 100 admissions) were calculated for frequently used drug groups (by ATC level 1 and 3) and APIs (ATC level 5) for all GA groups and regions. Formulations that consist only one vitamin (e.g. vitamin D, phytomenadione) were analyzed separately. All multivitamin products (enteral and parenteral formulations) were analyzed as group named “multivitamins”. Different vitamins in multivitamin compositions (e.g. vitamin D) were not calculated separately.

### Statistical analysis

Statistical software R (version 3.1.1.) was used for analyses. The impact of GA and region on drug use was analyzed using uni-and multivariate logistic regression analyses, where the East region and extremely preterms were reference groups. The primary analyses included the six most commonly used therapeutic groups (ATC level 1).

Additional multivariate analyses included twelve commonly used subgroups (ATC level 3). Vitamins, minerals and probiotics were excluded from additional analyses as these groups have the smallest evidence base for use.

## RESULTS

### Study population

Out of 31 invited countries, 21 participated (Austria, Belgium, Bulgaria, England, Estonia, France, Greece, Hungary, Ireland, Italy, Latvia, Lithuania, Malta, Netherlands, Portugal, Romania, Slovenia, Spain and 3 non-EU countries: Norway, Serbia, Switzerland) with 89 units from 73 hospitals; both relatively evenly distributed between regions (Figure 1).

**Figure 1.**
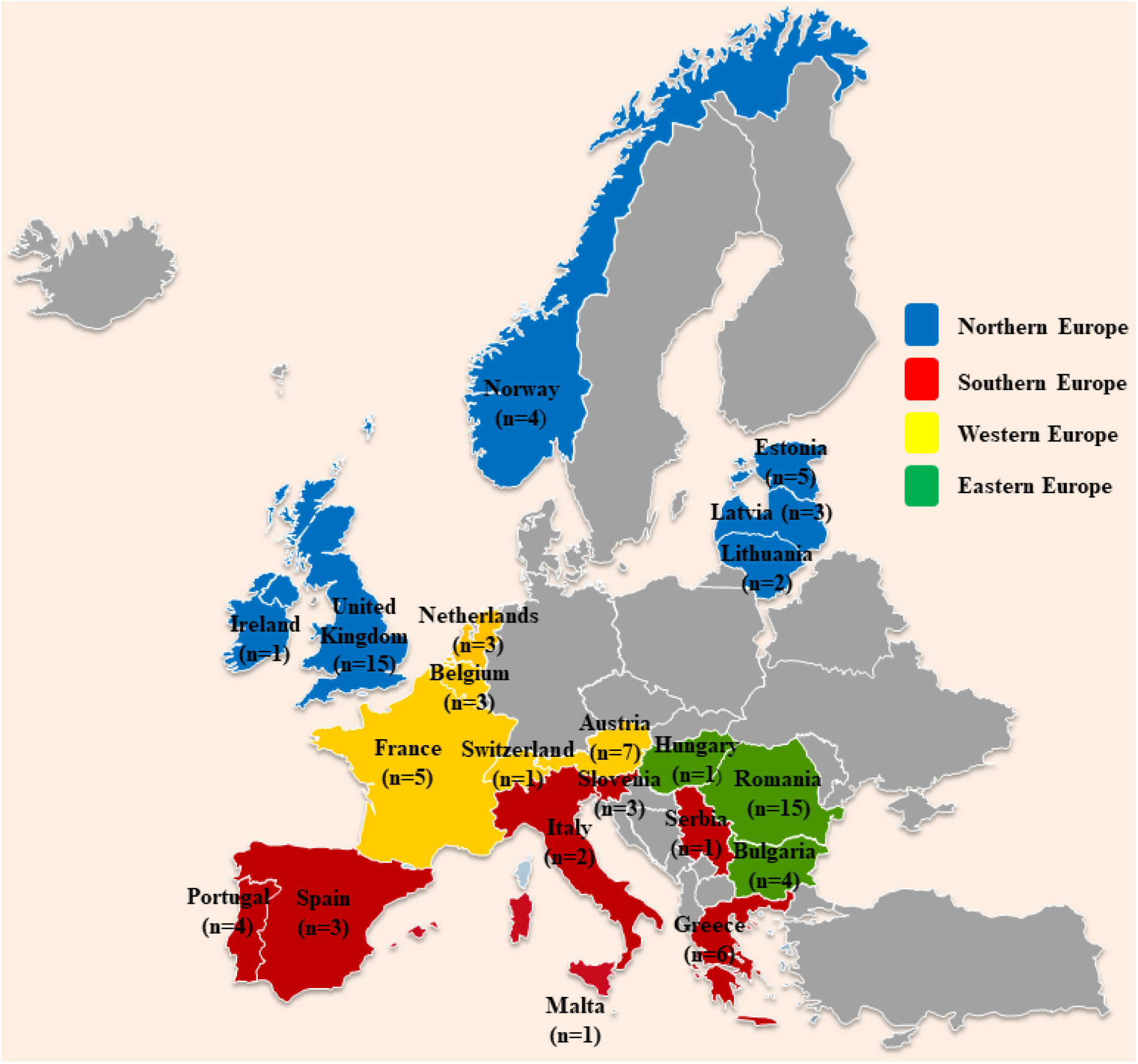
Participating countries by European region (shown in different colors). Number of neonatal units participating from each country is shown in parentheses.

Data of 726 patients of whom almost two thirds were preterm neonates (n=477, 65.7%), were collected. The proportional distribution of neonates based on GA varied significantly between regions, with the highest representation of extremely preterms in the West (21%) and of term neonates in the East (52.4%) (Table 1).

**Table 1.** Distribution of participants based on geographic region

|  | Geographic region |  |  |  | Total | p value |
| --- | --- | --- | --- | --- | --- | --- |
|  | East | North | South | West |  |  |
| No (%) of participating hospitals | 16 (21.9) | 26 (35.6) | 18 (24.7) | 13 (17.8) | 73 (100) | NS |
| No (%) of participating units | 20 (22.5) | 30 (34) | 20 (22.5) | 19 (21) | 89 (100) | NS |
| No (%) of neonates receiving any prescription during the study period | 185 (25.5) | 274 (37.7) | 143 (19.7) | 124 (17.1) | 726 (100) | **p<0.001** |
| Distribution of children in gestational age groups, n (%): |  |  |  |  |  | **p<0.001<sup>1</sup>** |
| Extremely preterm | 8 (4.3) | 40 (14.6) | 10 (7.0) | 26 (21.0) | 84 (11.6) |  |
| Very preterm | 29 (15.7) | 69 (25.2) | 28 (19.6) | 36 (29.0) | 162 (22.3) |  |
| Late preterm | 51 (27.6) | 87 (31.8) | 54 (37.8) | 39 (31.5) | 231 (31.8) |  |
| Term neonates | 97 (52.4) | 78 (28.5) | 51 (35.7) | 23 (18.5) | 249 (34.3) |  |
| Distribution of neonates based on neonatal units' level of care, n (%) |  |  |  |  |  | **p<0.001<sup>2</sup>** |
| Level 1 | 7 (3.8) | 0 (0.0) | 0 (0.0) | 4 (3.2) | 11 (1.5) |  |
| Level 2 | 54 (29.2) | 61 (22.3) | 18 (12.6) | 11 (8.9) | 144 (19.8) |  |
| Level 3 | 124 (67.0) | 213 (77.7) | 125 (87.4) | 109 (87.9) | 571 (78.7) |  |
| Distribution of neonates based on hospital teaching status, n (%) |  |  |  |  |  | **p<0.001<sup>3</sup>** |
| No | 47 (25.4) | 34 (12.4) | 7 (4.9) | 33 (26.6) | 121 (16.7) |  |
| Yes | 138 (74.6) | 240 (87.6) | 136 (95.1) | 91 (73.4) | 605 (83.3) |  |

P-values from Chi-squared test; NS- not significant at $\alpha = 0.05$ . Percentages may not sum to 100 because of rounding.
<sup>1</sup> significant differences in neonates' distribution into GA groups between regions e.g. the proportion of term neonates in the Eastern region and the proportion of extremely preterm neonates in the Western region are significantly higher compared to other regions; <sup>2</sup> significant differences in the distribution of department levels between regions – the proportion of Level 1 and Level 2 hospitals was the highest in the Eastern region; <sup>3</sup> significant differences in the distribution of participating units' teaching statuses between regions – the proportion of teaching hospitals was higher in the Southern and Northern region compared to the Eastern and Western region.

The distribution of neonates based on level of maturity is shown in Table 2.

**Table 2.** Distribution of study population according to level of maturity

|  | **Extremely**<br>**preterm**<br>**neonates** | **Very**<br>**preterm**<br>**neonates** | **Late**<br>**preterm**<br>**neonates** | **Term**<br>**neonates** | **Total** | ***p* value** |
| --- | --- | --- | --- | --- | --- | --- |
| No of patients admitted | 84 | 162 | 231 | 249 | 726 | <0.001 |
| Sex, male<sup>1</sup> | 47 | 93 | 142 | 129 | 411 | NS |
| Proportional distribution (%)<br>of neonates by region |  |  |  |  |  | <0.001 |
| East | 9.5 | 17.9 | 22.1 | 39 | 25.5 |  |
| North | 47.6 | 42.6 | 37.7 | 31.3 | 37.7 |  |
| South | 11.9 | 17.3 | 23.4 | 20.5 | 19.7 |  |
| West | 31 | 22.2 | 16.9 | 9.2 | 17.1 |  |
| Birth weight (grams) median<br>(IQR) | 830<br>(686–940) | 1340<br>(1140–1580) | 1980<br>(1718–2310) | 3340<br>(2960–3680) | 1993<br>(1356–3006) | <0.001 |
| Apgar score at 1 minute<sup>2</sup><br>mean ± SD | 5.3±2.4 | 6.3±2.4 | 7.4±1.9 | 7.9±1.9 | 7.1±2.3 | <0.001 |
| GA weeks<sup>3</sup> median (IQR) | 26 (25–27) | 30 (28–31) | 34 (33–35) | 39 (38–40) | 34 (30–38) | <0.001 |
| Postnatal age on study day<br>morning (days) median (IQR) | 15<br>(7–21.5) | 13<br>(7–20) | 10<br>(5–18) | 4<br>(1–15) | 10<br>(3–18) | <0.001 |

<sup>1</sup> Data not available for 1 neonate; <sup>2</sup> Data not available for 22 neonates;
<sup>3</sup> Full gestation weeks; P-values from Chi-squared test; NS- not significant at $\alpha = 0.05$

### Prescriptions

In total 2173 prescriptions with median number of 2 prescriptions per neonate [interquartile range (IQR) 1–4] were registered.

The most commonly prescribed drug groups based on ATC level 1 were drugs for alimentary tract and metabolism (A, 31%), systemic anti-infectives (J, 26%), drugs for blood and blood-forming organs (B, 24%), nervous (N, 11%), respiratory (R, 3%) and cardiovascular system (C, 3%). These six groups accounted for 98% of all prescriptions and were included into further analysis. The distribution of prescriptions is presented in Table 3.

**Table 3.** Distribution of registered prescriptions

|  | Gestational age group |  |  |  |  | Geographic region |  |  |  |  | Total |
| --- | --- | --- | --- | --- | --- | --- | --- | --- | --- | --- | --- |
|  | Extremely preterm | Very preterm | Late preterm | Term neonates | *p* value<sup>†</sup> | East | North | South | West | *p* value<sup>†</sup> |  |
| Total number of prescriptions (rate*) | 394 (469) | 603 (372) | 612 (265) | 564 (227) | **<0.001** | 503 (272) | 741 (270) | 404 (283) | 525 (423) | **<0.001** | 2173 (299) |
| No of prescriptions per neonate, median (IQR) | 4 (3–6) | 3 (2–5) | 2 (1–3) | 2 (1–3) |  | 2 (1–4) | 2 (2–3) | 2 (1–4) | 4 (2–6) |  | 2 (1–4) |
| Distribution of prescriptions based on ATC level 1, No (rate*) |  |  |  |  |  |  |  |  |  |  |  |
| Alimentary tract and metabolism | 109 (130) | 199 (123) | 240 (104) | 128 (51) | **<0.001** | 148 (80) | 235 (86) | 100 (70) | 193 (156) | **<0.001** | 676 (93) |
| Anti-infectives | 82 (98) | 128 (79) | 153 (66) | 209 (84) | NS | 134 (72) | 223 (81) | 147 (103) | 68 (55) | **0.001** | 572 (79) |
| Blood and blood-forming organs | 98 (117) | 159 (98) | 144 (62) | 117 (47) | **<0.001** | 152 (82) | 140 (51) | 65 (45) | 161 (130) | **<0.001** | 518 (71) |
| Nervous system | 70 (83) | 75 (46) | 37 (16) | 48 (19) | **<0.001** | 20 (11) | 87 (32) | 51 (36) | 72 (58) | **<0.001** | 230 (32) |
| Respiratory system | 9 (11) | 31 (19) | 19 (8) | 15 (6) | **0.006** | 29 (16) | 14 (5) | 21 (15) | 10 (8) | **0.006** | 74 (10) |
| Cardiovascular system | 17 (20) | 9 (6) | 7 (3) | 23 (9) | **0.003** | 9 (5) | 30 (11) | 6 (4) | 11 (9) | NS | 56 (8) |
| Other groups | 9 (11) | 2 (1) | 12 (5) | 24 (10) | **<0.001** | 11 (6) | 12 (4) | 14 (10) | 10 (8) | NS | 47 (6) |

\*Rate – number of prescriptions per 100 admissions; <sup>†</sup>p-values from Quasi-Poisson model; NS- not significant at $\alpha = 0.05$

The most commonly used drugs based on ATC level 5 were multivitamins, followed by vitamin D and caffeine. Among the ten most commonly used APIs were three systemic anti-infectives – gentamicin, ampicillin and benzylpenicillin (Table 4).

**Table 4.** Prescription rates (prescriptions per 100 admissions) of most frequently used drugs (based on ATC level 5) according to GA and geographic region

|  | Total | Gestational age group |  |  |  |  | Geographic region |  |  |  |
| --- | --- | --- | --- | --- | --- | --- | --- | --- | --- | --- |
|  |  | Extremely preterm | Very preterm | Late preterm | Term |  | East | North | South | West |
| Multivitamins | 33 | **43\*** | **46\*** | **45\*** | 10 |  | 5 | **33\*** | **31\*** | **74\*** |
| Vitamin D | 19 | 24 | 19 | **23\*** | 14 |  | **23\*** | 18 | 13 | 23 |
| Caffeine | 19 | **60\*** | **43\*** | 6 | 1 |  | 2 | 22 | 20 | **37\*** |
| Gentamicin | 18 | 19 | 12 | 16 | **24\*** |  | 11 | **25\*** | **23\*** | 7 |
| Amino acids for parenteral nutrition | 18 | **31\*** | **25\*** | **19\*** | 7 |  | 19 | 12 | 20 | 24 |
| Phytomenadione | 13 | 6 | 9 | 3 | **26\*** |  | **30\*** | 7 | 1 | 13 |
| Ampicillin (/+sulbactam) | 12 | 5 | 10 | **19\*** | 11 |  | **20\*** | 3 | **26\*** | 6 |
| Benzylpenicillin | 12 | 12 | 10 | 8 | **15\*** |  | 3 | **24\*** | 2 | 8 |
| Fat emulsion for parenteral nutrition | 11 | 24 | 18 | 10 | 3 |  | 7 | 7 | 8 | **30\*** |
| Probiotics | 8 | 19 | 13 | 7 | 2 |  | 5 | 6 | 3 | 23 |
| Iron | 8 | 17 | 17 | 7 | 1 |  | 9 | 3 | 6 | 20 |
| Heparin | 7 | 19 | 7 | 6 | 4 |  | 2 | 9 | 4 | 15 |
| Folic acid | 7 | 7 | 13 | 10 | 1 |  | 5 | 9 | 4 | 7 |
| Aminophylline | 6 | 7 | 14 | 6 | 0 |  | 14 | 1 | 10 | 0 |
| Vancomycin | 5 | 12 | 6 | 3 | 4 |  | 4 | 4 | 9 | 7 |

\*The three most commonly used drugs in each group.
This table does not contain drugs which were not included among 15 most commonly used drugs in general (e.g. fluconazole and nystatin were common among extremely preterm neonates, but not in other GA groups).

The forty most commonly prescribed drugs are listed in Table 5.

**Table 5.**
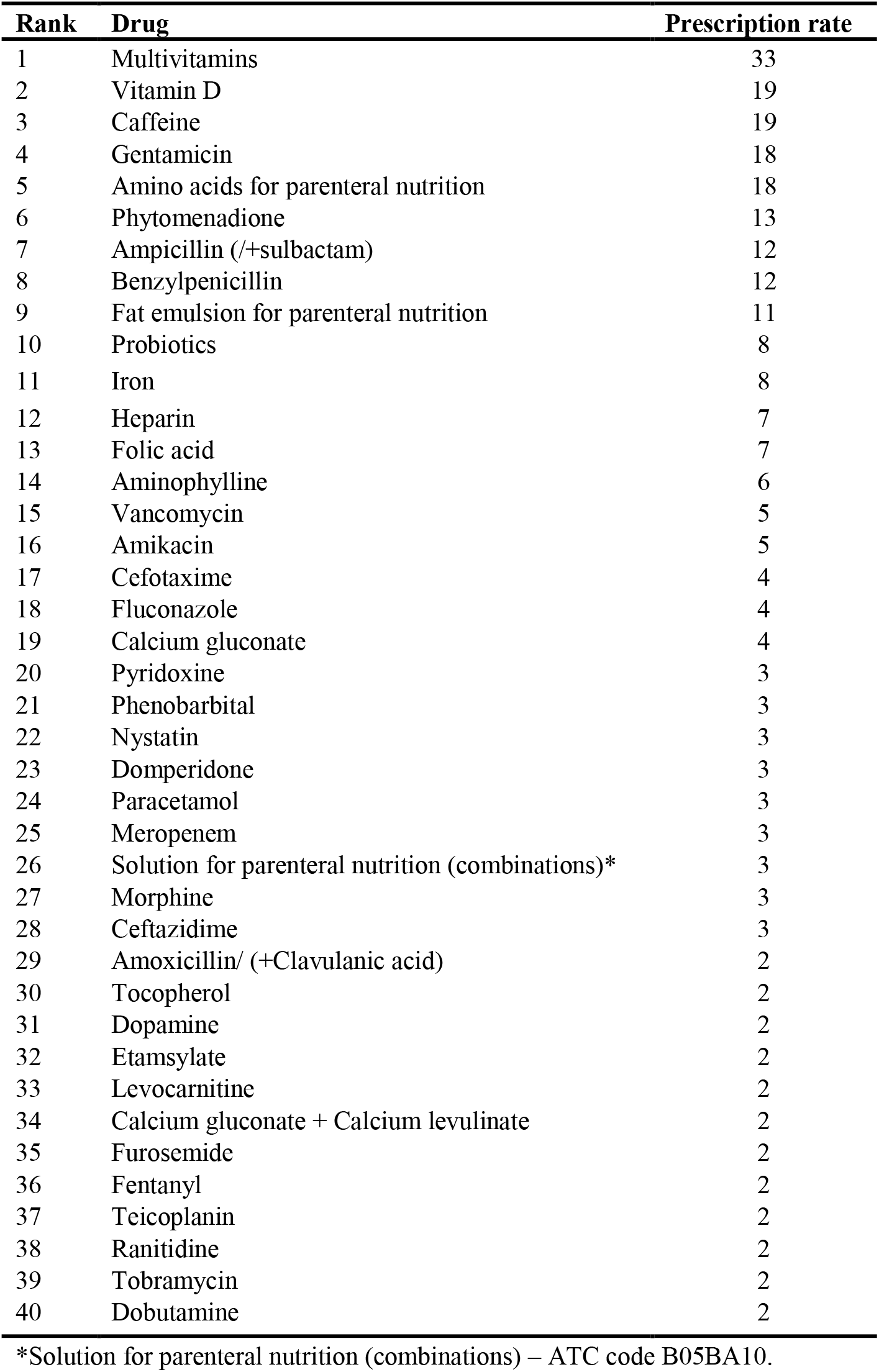
Drugs most commonly used in neonatal units

| **Rank** | **Drug** | **Prescription rate** |
| --- | --- | --- |
| 1 | Multivitamins | 33 |
| 2 | Vitamin D | 19 |
| 3 | Caffeine | 19 |
| 4 | Gentamicin | 18 |
| 5 | Amino acids for parenteral nutrition | 18 |
| 6 | Phytomenadione | 13 |
| 7 | Ampicillin (/+sulbactam) | 12 |
| 8 | Benzylpenicillin | 12 |
| 9 | Fat emulsion for parenteral nutrition | 11 |
| 10 | Probiotics | 8 |
| 11 | Iron | 8 |
| 12 | Heparin | 7 |
| 13 | Folic acid | 7 |
| 14 | Aminophylline | 6 |
| 15 | Vancomycin | 5 |
| 16 | Amikacin | 5 |
| 17 | Cefotaxime | 4 |
| 18 | Fluconazole | 4 |
| 19 | Calcium gluconate | 4 |
| 20 | Pyridoxine | 3 |
| 21 | Phenobarbital | 3 |
| 22 | Nystatin | 3 |
| 23 | Domperidone | 3 |
| 24 | Paracetamol | 3 |
| 25 | Meropenem | 3 |
| 26 | Solution for parenteral nutrition (combinations)* | 3 |
| 27 | Morphine | 3 |
| 28 | Ceftazidime | 3 |
| 29 | Amoxicillin/ (+Clavulanic acid) | 2 |
| 30 | Tocopherol | 2 |
| 31 | Dopamine | 2 |
| 32 | Etamsylate | 2 |
| 33 | Levocarnitine | 2 |
| 34 | Calcium gluconate + Calcium levulinate | 2 |
| 35 | Furosemide | 2 |
| 36 | Fentanyl | 2 |
| 37 | Teicoplanin | 2 |
| 38 | Ranitidine | 2 |
| 39 | Tobramycin | 2 |
| 40 | Dobutamine | 2 |

\*Solution for parenteral nutrition (combinations) – ATC code B05BA10.

### Drug use and GA

Compared to extremely preterms there was a significant downwards trend in drug use with increasing GA – the prescription rate among extremely preterms was two times higher than term neonates (469/100 vs 227/100, respectively) (Table 3).

The list of commonly used APIs varied between GA groups. Caffeine was the most commonly used medicine among extremely preterms and frequently used among very preterms, however, it was rarely used among late preterm and term neonates as expected. Multivitamins were more commonly used among preterms compared to term neonates. Vitamin D was used with similar frequency among preterms (Table 4), but slightly less among term neonates.

Anti-infectives’ prescription rate was the highest among extremely preterms, who received most commonly gentamicin (19/100), fluconazole (18/100), nystatin (13/100), vancomycin (12/100) and benzylpenicillin (12/100). The use of fluconazole and nystatin was lower among very preterms (5/100 and 4/100, respectively) and rare among late preterm and term neonates.

In univariate analysis, significant GA-related variations were observed in all six commonly used ATC level 1 groups. As shown in Table 6, extremely preterms had higher odds of receiving nervous system and cardiovascular drugs compared to other groups, drugs for blood and blood-forming organs and alimentary system compared to late preterm and term neonates; anti-infectives compared to late preterms and respiratory system drugs compared to term neonates.

**Table 6.** Odd ratios (OR) of receiving commonly used ATC level 1 drug groups, based on univariate logistic regression analysis

|  | Gestational age group |  |  |  | Geographic region |  |  |  |
| --- | --- | --- | --- | --- | --- | --- | --- | --- |
|  | Extremely preterm | Very preterm | Late preterm | Term neonates | East | North | South | West |
| Alimentary tract and metabolism (A) | 1 | 0.86<br>(0.46–1.62) | **0.55**<br>**(0.3–0.98)\*** | **0.14**<br>**(0.08–0.25)\*** | 1 | 1.39<br>(0.95–2.02) | 0.95<br>(0.61–1.47) | **5.14**<br>**(2.94–8.99)\*** |
| Anti-infectives (J) | 1 | 0.59<br>(0.35–1.01) | **0.55**<br>**(0.33–0.91)\*** | 0.85<br>(0.52–1.39) | 1 | 1.01<br>(0.7–1.48) | **1.57**<br>**(1.01–2.43)\*** | 0.66<br>(0.41–1.06) |
| Blood and blood-forming organs (B) | 1 | 0.6<br>(0.35–1.06) | **0.29**<br>**(0.17–0.5)\*** | **0.26**<br>**(0.15–0.45)\*** | 1 | **0.4**<br>**(0.27–0.58)\*** | **0.31**<br>**(0.19–0.48)\*** | 1.06<br>(0.66–1.68) |
| Nervous system (N) | 1 | **0.35**<br>**(0.2–0.61)\*** | **0.07**<br>**(0.04–0.12)\*** | **0.07**<br>**(0.04–0.12)\*** | 1 | **3.66**<br>**(2.08–6.44)\*** | **3.97**<br>**(2.14–7.36)\*** | **9.26**<br>**(5.03–17.06)\*** |
| Respiratory system (R) | 1 | 1.74<br>(0.78–3.88) | 0.66<br>(0.28–1.55) | **0.31**<br>**(0.12–0.82)\*** | 1 | **0.19**<br>**(0.09–0.41)\*** | 0.76<br>(0.4–1.44) | **0.44**<br>**(0.2–0.97)\*** |
| Cardiovascular system (C) | 1 | **0.31**<br>**(0.12–0.8)\*** | **0.16**<br>**(0.06–0.44)\*** | **0.38**<br>**(0.17–0.86)\*** | 1 | 2<br>(0.83–4.84) | 0.92<br>(0.29–2.97) | 1.99<br>(0.72–5.49) |

\* significant at $\alpha = 0.05$

In multivariate analysis (Table 7) after adjusting for region, GA remained significant in the prescription of the following ATC level 1 categories in comparison to extremely preterms: anti-infectives and cardiovascular agents were less frequently used in very and late preterms; drugs for blood and blood-forming organs in late preterms and terms; respiratory and alimentary drugs in term neonates and nervous system agents in all other GA groups. Based on ATC level 3 other antibacterials (J01X) and iron were less frequently used in late preterm and term babies; opioids in very and late preterms; and psychostimulants, systemic antimycotics, cardiac stimulants and antithrombotic agents in all other GA groups compared to extremely preterms while phytomenadione was used more frequently in term babies.

**Table 7.** Multivariate logistic regression analysis. The odds (OR) of neonates receiving commonly used ATC groups†

|  |  | Gestational age group |  |  |  | Geographic region |  |  |  |
| --- | --- | --- | --- | --- | --- | --- | --- | --- | --- |
| ATC level 1 | ATC level 3<br>(frequently used<br>active ingredients,<br>arranged by<br>frequency of use) | Extremely<br>preterm<br>(reference)<br>OR (95% CI) | Very<br>preterm<br>OR (95% CI) | Late<br>preterm<br>OR (95% CI) | Term<br>OR (95% CI) | East<br>(reference)<br>OR (95% CI) | North<br>OR (95% CI) | South<br>OR (95% CI) | West<br>OR (95% CI) |
| **Alimentary tract**<br>**and metabolism**<br>**(A)** | (Multivitamins,<br>vitamin D,<br>antidiarrheal<br>microorganisms) | 1 | 0.95<br>(0.49–1.81) | 0.63<br>(0.34–1.16) | **0.16**<br>**(0.09–0.3)\*** | 1 | 0.93<br>(0.62–1.41) | 0.71<br>(0.44–1.14) | **3.28**<br>**(1.81–5.94)\*** |
| **Anti-infectives**<br>**(J)** |  | 1 | **0.55**<br>**(0.32–0.95)\*** | **0.48**<br>**(0.29–0.81)\*** | 0.74<br>(0.44–1.25) | 1 | 1.03<br>(0.7–1.52) | **1.65**<br>**(1.06–2.58)\*** | 0.66<br>(0.4–1.09) |
|  | **Penicillins (J01C)**<br>(Ampicillin,<br>Benzylpenicillin,<br>Amoxicillin) | 1 | 1.18<br>(0.62–2.23) | 1.47<br>(0.8–2.68) | 1.75<br>(0.95–3.2) | 1 | **1.68**<br>**(1.09–2.6)\*** | 1.35<br>(0.82–2.23) | 0.99<br>(0.56–1.74) |
|  | **Aminoglycosides**<br>**(J01G)**<br>(Gentamicin,<br>Amikacin) | 1 | 0.86<br>(0.44–1.7) | 1.02<br>(0.54–1.93) | 1.58<br>(0.85–2.96) | 1 | **2.17**<br>**(1.35–3.5)\*** | **2.4**<br>**(1.42–4.07)\*** | 0.67<br>(0.33–1.36) |
|  | **Other beta-lactams**<br>**(J01D)**<br>(Cefotaxime,<br>Meropenem,<br>Ceftazidime) | 1 | 0.79<br>(0.36–1.72) | 0.46<br>(0.21–1) | 0.73<br>(0.35–1.54) | 1 | 0.68<br>(0.37–1.25) | 1.69<br>(0.92–3.09) | 0.49<br>(0.21–1.14) |
|  | **Other**<br>**antibacterials**<br>**(J01X)**<br>(Vancomycin,<br>Teicoplanin,<br>Metronidazole) | 1 | 0.63<br>(0.29–1.34) | **0.21**<br>**(0.09–0.5)\*** | **0.23**<br>**(0.1–0.54)\*** | 1 | 0.66<br>(0.3–1.47) | **2.47**<br>**(1.16–5.26)\*** | 0.82<br>(0.33–2.05) |
|  | **Systemic**<br>**antimycotics**<br>**(J02A)**<br>(Fluconazole) | 1 | **0.13**<br>**(0.04–0.36)\*** | **0.03**<br>**(0.01–0.13)\*** | **0.01**<br>**(0–0.05)\*** | 1 | **0.07**<br>**(0.02–0.23)\*** | **0.12**<br>**(0.03–0.49)\*** | **0.13**<br>**(0.04–0.41)\*** |

Table 7. Continued
|  |  | Gestational age group |  |  |  | Geographic region |  |  |  |
| --- | --- | --- | --- | --- | --- | --- | --- | --- | --- |
| ATC level 1 | ATC level 3<br>(frequently used<br>active ingredients,<br>arranged by<br>frequency of use) | Extremely<br>preterm<br>(reference)<br>OR (95% CI) | Very preterm<br>OR (95% CI) | Late preterm<br>OR (95% CI) | Term<br>OR (95% CI) | East<br>(reference)<br>OR (95% CI) | North<br>OR (95% CI) | South<br>OR (95% CI) | West<br>OR (95% CI) |
| Blood and blood-forming organs<br>(B) |  | 1 | 0.58<br>(0.33–1.03) | **0.26**<br>(0.15–0.46)* | **0.2**<br>(0.11–0.35)* | 1 | **0.28**<br>(0.19–0.43)* | **0.26**<br>(0.16–0.41)* | 0.7<br>(0.43–1.16) |
|  | **Vitamin K and**<br>**other hemostatics**<br>**(B02B)**<br>(Phytomenadione,<br>Etamsylate) | 1 | 1.01<br>(0.38–2.68) | 0.5<br>(0.18–1.39) | **2.94**<br>(1.21–7.13)* | 1 | **0.16**<br>(0.09–0.29)* | **0.01**<br>(0–0.11)* | **0.4**<br>(0.21–0.77)* |
|  | **Iron (B03A)** | 1 | 1<br>(0.47–2.11) | **0.38**<br>(0.17–0.84)* | **0.03**<br>(0.01–0.16)* | 1 | **0.18**<br>(0.08–0.44)* | 0.47<br>(0.2–1.12) | 1.24<br>(0.6–2.55) |
|  | **Antithrombotic**<br>**agents (B01A)**<br>(Heparin) | 1 | **0.39**<br>(0.18–0.86)* | **0.3**<br>(0.13–0.65)* | **0.26**<br>(0.11–0.61)* | 1 | **3.32**<br>(1.11–9.94)* | 1.83<br>(0.5–6.67) | **6.35**<br>(2.05–20)* |
| Nervous system<br>(N) |  | 1 | **0.37**<br>(0.21–0.66)* | **0.07**<br>(0.04–0.13)* | **0.09**<br>(0.05–0.16)* | 1 | **2.64**<br>(1.44–4.83)* | **4.17**<br>(2.16–8.07)* | **6.67**<br>(3.43–13)* |
|  | **Psychostimulants**<br>**(N06B)** (Caffeine) | 1 | **0.52**<br>(0.29–0.93)* | **0.04**<br>(0.02–0.09)* | **0.01**<br>(0–0.03)* | 1 | **10.55**<br>(3.1–36)* | **18.75**<br>(5.16–68)* | **23.7**<br>(6.7–84)* |
|  | **Opioids (N02A)**<br>(Morphine,<br>Fentanyl) | 1 | **0.2**<br>(0.06–0.68)* | **0.23**<br>(0.08–0.65)* | 0.44<br>(0.17–1.14) | 1 | 3.97<br>(0.86–18) | **4.99**<br>(1.01–25)* | **6.46**<br>(1.31–32)* |
| Respiratory<br>system (R) |  | 1 | 1.4<br>(0.6–3.24) | 0.42<br>(0.17–1.03) | **0.15**<br>(0.05–0.42)* | 1 | **0.11**<br>(0.05–0.26)* | 0.59<br>(0.3–1.16) | **0.23**<br>(0.1–0.53)* |
|  | **Other systemic**<br>**drugs for**<br>**obstructive airway**<br>**disease (R03D)**<br>(Aminophylline) | 1 | 1.75<br>(0.63–4.87) | 0.48<br>(0.16–1.39) | 0<br>(0–Inf) | 1 | **0.02**<br>(0–0.09)* | **0.41**<br>(0.19–0.88)* | **0.1**<br>(0.03–0.28)* |
| Cardiovascular<br>system (C) |  | 1 | **0.33**<br>(0.13–0.86)* | **0.18**<br>(0.06–0.51)* | 0.48<br>(0.2–1.11) | 1 | 1.91<br>(0.76–4.76) | 0.98<br>(0.3–3.18) | 1.78<br>(0.61–5.21) |
|  | **Cardiac stimulants**<br>**(C01C)** (Dopamine,<br>Dobutamine) | 1 | **0.06**<br>(0.01–0.47)* | **0.17**<br>(0.05–0.58)* | **0.2**<br>(0.06–0.66)* | 1 | 1.78<br>(0.45–6.95) | 1.23<br>(0.24–6.36) | 0.64<br>(0.1–4.24) |

† adjusted for GA and geographic region; \* significant at $\alpha = 0.05$

### Drug use and region

The West region stood out with a significantly higher prescription rate (423/100) compared to others, where the rate was similar (270/100 North, 272/100 East, 283/100 South) (Table 3).

The list of most frequently used drugs varied between regions – while in the North, South and West multivitamins dominated, phytomenadione was the most commonly used in the East (Table 4). Although multivitamins were the most commonly used drugs in three regions, we observed high variability in composition of multivitamin products used between countries and hospitals.

Univariate analysis revealed significant geographic variations in all the most commonly used ATC groups except cardiovascular drugs. Compared to neonates in the East those in the West received more commonly medicines for alimentary tract, in the South anti-infectives and in all other regions medicines for nervous system. Neonates in the West and North received fewer drugs for respiratory system and in the North and South for blood and blood-forming organs compared to the East. The high use of drugs for blood and blood-forming organs in the East was mainly due to phytomenadione use in term neonates.

In multivariate analysis when adjusting for GA, region remained significant in all above-mentioned drug groups (Table 7). Based on ATC level 3 in comparison to the East psychostimulants (caffeine) were more commonly used in all other regions, penicillins in the North, aminoglycosides in the North and South, other antibacterials (e.g. vancomycin) in the South, antithrombotic agents in the North and West and opioids in the South and West regions. The use of iron supplements was lower in the North and the use of systemic antimycotics, phytomenadione and other systemic drugs for obstructive airway disease were lower in all regions compared to the East (Table 7).

## DISCUSSION

To the best of our knowledge this is the largest multi-country survey of drug use covering neonatal units across Europe and neonates of all GA groups. We have made the following observations: first, the most commonly used drugs in European neonatal units were multivitamins and the composition of these products was highly variable. Second, there is a significant variation in drug use depending on GA as expected but also unexpectedly depending on geographical region. While GA-based variations could be explained by different underlying conditions, geographic variations could refer to the lack of standardized evidence-based guidelines for treatment. Variations in anti-infectives’ use may reflect different patterns of infections or resistance as described elsewhere [15]. Third, although neonates receive many drugs during hospitalization, basically no specific medicines are available for treatment or prevention of conditions affecting especially neonates and directly related to their mortality like respiratory distress syndrome, intraventricular hemorrhage (IVH), perinatal asphyxia and/or necrotizing enterocolitis (NEC) [16]. This suggests the urgent need to understand the underlying mechanisms of these conditions followed by the development of new treatment options (including drugs).

We demonstrated that drugs for alimentary tract were by far most commonly used mainly due to high prescription rate of vitamins and probiotics similarly shown in previous studies [1]. High variability in multivitamin products composition could be explained by the lack of international guidelines despite the availability of national recommendations suggesting multivitamins for babies born at GA <34 weeks. However, there is limited evidence of required vitamin quantities [17,18].

Consistent with other studies, systemic anti-infectives were commonly used with four antibiotics (gentamicin, ampicillin, benzylpenicillin, vancomycin) belonging to the 15 most common APIs [19,20]. This is not surprising as many neonates in neonatal units have either risk factors for getting infections or confirmed bacterial infections [21]. High use of penicillins, aminoglycosides and other beta-lactams in neonatal units has also been shown in previous studies [22]. In multivariate analysis anti-infectives were less commonly given to very and late preterms than to other groups. We suggest that reasons for that are higher systemic antibacterials’ use in extremely preterm group due to higher rate of infection and that infections are likely one of the most frequent cause of hospitalization of term neonates.

Similarly to previous studies we demonstrate a negative correlation between GA and number of drugs per patient [1,19,23,24]. We found that extremely preterms received drugs for alimentary tract and metabolism (e.g. multivitamins, vitamin D, amino acids) given likely as supplementation recommended by learned societies to support growth and development significantly more commonly compared to term neonates [25,26].

In contrast to our hypothesis, we observed significant regional differences in drug use even after adjustment for GA. We believe that geographical differences in anti-infectives’ use are partly explained by differences in microbial resistance rate – the microorganisms causing infection in the South are generally more resistant compared to those in other regions [27]. However, differences in medical practices cannot be excluded. Regional variations in systemic antimycotics’ use with higher prescription rate in the East compared to other regions are likely driven by the differences in local guidelines as shown in a pan-European study [28]. It is important to emphasize that systemic antimycotics are mainly used for prophylaxis since invasive fungal infections in neonates have become extremely rare [29]. Similar to ARPEC study we showed higher penicillins’ use in the North, higher aminoglycosides’ use in the South and North, and higher use of other antibacterials’ (e.g. vancomycin) in the South [30].

Regional differences in other drugs’ use are more complex to explain. For example, less frequent use of caffeine or more frequent use of phytomenadione and aminophylline in the East, lower use of iron in the North compared to other regions or more frequent use of heparin in the North and West than in the East and South or greater use of opioids in the South and West compared to the East and North need further investigating. While studies have shown beneficial effect of caffeine in reducing bronchopulmonary dysplasia in preterms with apnea the data on using bronchodilators (e.g. aminophylline) are less convincing according to the published literature [31,32]. Although opioids are routinely used in neonatal units for analgesia and sedation, their possible long-term negative impact is unclear due to conflicting results concerning neurodevelopmental outcome [33,34]. We believe that these regional differences are partly explained by the lack of reliable studies (e.g. bronchodilators) or failure to implement results of randomized controlled trials (RCTs) into clinical practice (e.g. caffeine) suggesting that further efforts should be focused to both directions.

Some limitations need to be noted. The voluntary participation prevented including the data from whole Europe proportionally and led to missing data from some large countries (e.g. Germany). Due to inability to randomly select participants, some small countries (e.g. Estonia) were covered almost completely, while large countries were represented with a few areas. As most participants were from teaching hospitals and level III neonatal units, it was impossible to detect the impact of teaching status and departments’ level on the prescription pattern. Neonates were not divided equally between GA groups and regions which could also impact the effect of analyzed factors. In this study moderately preterm neonates (32-34 weeks) were considered “late preterms” (32-36 weeks), which could affect some results as these newborns may behave as very preterm infants or as late preterms. Also, postnatal age which was not taken into account could affect results of some medicines’ administration (e.g. phytomenadione). The selection of study methodology can affect the results of this study – for medicines that are commonly used (e.g. antibiotics like ampicillin, gentamycin or phytomenadione) one day PPS will describe their consumption sufficiently, however, for agents that are rarely used longer study period will capture more APIs. Although vitamin D was among 3 most commonly used drugs in Europe, we did not add it into regional analysis as hospitalized neonates could receive it from different sources e.g. multivitamin compositions or breast milk fortifiers. The data about last-mentioned products was not collected according to the ESNEE study protocol and multivitamin compositions were not viewed separately, which makes it impossible to compare the vitamin D prescription in different regions precisely. Nevertheless, we believe that none of these factors prevented us drawing adequate conclusions in describing drug use across Europe and analyzing regional differences.

This study indicates the need to prioritize research to the areas where there is the most urgent need to develop new drugs. It also suggests areas where clinical trials are needed to promote appropriate use of available drugs and to prevent potentially unnecessary prescriptions in a vulnerable population of neonates.

## CONCLUSION

This study highlights the influence of geographic region and GA on drug prescriptions in European neonatal units. While the impact of GA is explained by differences in maturation, requirements of regular nutritional supplementation and individualized care of preterm neonates, geographical differences with the exception of antimicrobials most likely indicate the lack of reliable RCTs and/or evidence-based guidelines for treatment of many important neonatal conditions. The presented data calls for greater collaboration between academia, basic scientists, practitioners, pharmaceutical industry and regulators in developing new drugs and support pan-European cooperation in facilitating neonatal drug development to improve the health and well-being of neonates.

## ABBREVIATIONS

API: active pharmaceutical ingredient
ATC: anatomical therapeutic chemical classification
ESNEE: European study of neonatal exposure to excipients
GA: gestational age
IQR: interquartile range
IVH: intraventricular haemorrhage
NEC: necrotizing enterocolitis
OR: odds ratio
PPS: point prevalence study
RCT: randomized control trial

## DECLARATIONS

### Ethics approval and consent to participate

Ethics Committee approval was obtained for participation in the study in compliance with national requirements. No consent for individual patients was sought, as the data were collected in routine clinical practice and anonymized before leaving the study sites.

### Consent for publication

Not applicable.

### Availability of data and material

The datasets used and/or analyzed during the current study are available from the corresponding author on reasonable request.

### Authors’ contributions

IM participated in study design, analyzed data and drafted the initial manuscript. GN participated in study design, data collection and revised the manuscript. JL participated in study design, data collection and drafting the manuscript. HVi performed statistical analysis and revised the manuscript. TM, HVa, MAT, AJN and JD participated in study design, data interpretation and revised the manuscript. IL participated in study design, coordinated and supervised data collection and interpretation, and drafting the manuscript. All authors read and approved the final manuscript.

## Acknowledgments

ESNEE consortium: Susan Graham (UK), Utpal Shah (UK), Hussain Mulla (UK), Hitesh Pandya (UK), James McElnay (UK), Jeff Millership (UK), Shirish Yakkundi (UK), Andre Rieutord (France), Thomas Storme (France), Pascal Vaconsin (France). All members of ESNEE designed the study and monitored data collection.

We thank all national contact persons who provided lists of neonatal units and helped run the study in their own country: Bernhard Resch (Austria), Pieter De Cock (Belgium), Nelly Jekova (Bulgaria), Elisabeth Iyore (Denmark), Pascal Vaconsin (France), Kosmas Sarafidis (Greece), Aranka Vegso (Hungary), Noreen O’Callaghan (Ireland), Rocco Agostino (Italy), Daiga Kviluna (Latvia), Rasa Tameliene (Lithuania), Rene F. Kornelisse (Netherlands), Dag Bratlid (Norway), Almerinda Pereira (Portugal), Maria Livia Ognean (Romania), Milica Bajcetic (Serbia), Darja Paro (Slovenia), Elizabeth Valls (Spain), Per Nydert (Sweden), Hans Ulrich Bucher (Switzerland), Maria Cordina (Malta).

We also thank local pharmacists for providing data on excipient content: Caroline Fonzo-Christe (Switzerland), Domenico Tarantino (Italy), Velina Grigorova (Bulgaria), Milica Bajcetic (Serbia), Elizabeth Valss (Spain), Claudine Milstein (France), Jennifer Duncan (England), Sabina Zalar (Slovenia), Per Gustaf Hartvig Honoré (Denmark).

